# A novel approach to modeling transcriptional heterogeneity identifies the oncogene candidate *CBX2* in invasive breast carcinoma

**DOI:** 10.1101/303396

**Authors:** Daniel G. Piqué, Cristina Montagna, John M. Greally, Jessica C. Mar

## Abstract

Oncogenes promote the development of and serve as therapeutic targets against subsets of cancers. Here, a new statistical approach that captures transcriptional heterogeneity in tumor and adjacent normal (i.e. tumor-free) mRNA expression profiles was developed to identify oncogene candidates that were overexpressed in a subset of breast tumors. Intronic DNA methylation was strongly associated with the overexpression of chromobox 2 (*CBX2*), an oncogene candidate that was identified using our method but not through prior analytical approaches. *CBX2* overexpression in breast tumors was associated with the upregulation of genes involved in cell cycle progression and is associated with poorer 5-year survival. The predicted function of *CBX2* was confirmed *in vitro* providing the first experimental evidence that *CBX2* promotes breast cancer cell growth. Modeling mRNA expression heterogeneity in tumors is a novel powerful approach with the potential to uncover therapeutic targets that benefit subsets of cancer patients.

## Introduction

Oncogenesis is driven by a complex and intricately controlled program of gene expression where oncogenes are the expressed genes that promote tumor development. The first set of oncogenes were discovered in retroviruses that incorporated human growth factors, such as *src*, into their viral genome^1–3^. The identification of amplified or mutated oncogenes in the tumors of certain cancer patients has led to the development of effective molecular therapeutic strategies that extend the life of these patients. For example, trastuzumab, an anti-Her2 antibody, extends overall lifespan for the approximately 20% of breast cancer patients whose often-aggressive tumors overexpress *ERBB2*, the gene that encodes the Her2 protein^4^. However, Her2-targeted therapies often result in treatment resistance, and thus additional therapeutic targets are required to adequately treat Her2^+^ breast cancer, among other subtypes.

Variability in the response of patients to current therapeutic strategies represents a major bottleneck to reducing cancer mortality rates globally. Understanding how tumor heterogeneity impacts the transcriptional regulatory programs that control oncogenesis is the key to addressing this issue and is currently what drives most programs in personalized medicine. The availability of genome-wide gene expression data from matched tumor and adjacent normal tissue of large patient populations provides a valuable resource for developing new approaches for identifying oncogenes that are likely to play pivotal roles in important clinical outcomes such as chemoresistance. Previous studies have identified survival-related biomarkers in ovarian cancer based on bimodal gene expression profiles detected in large datasets of tumors^5^. These studies recognize the limitations of the unimodal assumption made by many statistical tests and have taken advantage of the inherent heterogeneity in gene expression profiles to discover new subtypes.

Examples of methods that exploit heterogeneity between tumor and adjacent normal tissue include Cancer Outlier Profile Analysis (COPA)^6^ and mCOPA^7^, which are both used to detect gene fusions and tumor outliers. However, these kinds of approaches have two major limitations. First, most applications of mixture modeling for gene expression, with one exception^8^, have been developed using data derived from microarrays, which have a limited range of expression values, particularly for highly expressed genes, and unlike RNA-sequencing (RNA-seq), are limited for quantifying transcript levels at high resolution^9^. Second, tools developed for outlier detection from paired tumor-normal mRNA samples, such as cancer outlier profile analysis (COPA)^6,10^ and Profile Analysis using Clustering and Kurtosis (PACK)^11^, are sensitive to the proportion of samples that are distinguished as ‘outliers’^8^ and, in the case of COPA, require setting a tuning parameter. In addition, existing methods for outlier detection are designed to screen out individual tumor samples, rather than identify genes that reflect new patient subgroupings.

In this study, we developed a statistical approach termed *oncomix* to identify oncogene candidates in RNA-sequencing data. This approach detects oncogene candidates based on the presence of low expression in normal tissue and over-expression in a subset of patient tumors. Our approach capitalizes on the heterogeneity present in matched tumor and normal gene expression data to identify oncogene candidates and then segregate patients into interpretable subgroups based on their expression of the oncogene candidate. *Oncomix* is an unsupervised method where the size of the patient subgrouping is learned entirely from the data.

To demonstrate the utility of *oncomix*, we applied this approach to RNA-sequencing data from the breast cancer cohort of The Cancer Genome Atlas (TCGA) and identified a set of five high-confidence oncogene candidates (*CBX2, NELL2, EPYC, SLC24A2*, and *ZBED2*). To understand why these oncogene candidates were overexpressed in certain tumors, we developed predictive models using multiple molecular, genetic, and clinical variables from TCGA that highlighted potential regulators of oncogene candidate overexpression. Novel computational and experimental evidence suggest that chromobox 2 (CBX2), one of the oncogene candidates that we identified, is associated with poorer clinical outcomes and functions as a regulator of breast tumor cell growth. In this study, we demonstrate the value of modeling transcriptional heterogeneity using matched tumor and normal tissue to identify new oncogene candidates. Our results indicate that *CBX2* may serve as a driver of breast cancer and represent a novel therapeutic target.

## Results

### Deriving a new transcription-driven approach to discover oncogene candidates that are specific for subgroups of breast cancer patients

An oncogene candidate can be defined operationally as a gene that is highly expressed in a subset of tumor samples and has uniformly low expression in adjacent normal tissue. Our primary objective was to test whether such genes could be found in a cancer patient dataset. For this purpose, RNA-seq data from 110 breast cancer patients was selected from The Cancer Genome Atlas (TCGA). This population is predominantly represented by Caucasian females with infiltrating ductal carcinoma, that had both tumor and adjacent normal samples sequenced (Figure 1A). To ensure that the mixture models could be stably fit to the data, lowly-expressed genes were filtered (see Methods, Figure 1B). Two-component mixture models were fit to each transcript for both tumor and adjacent normal samples independently (Figure 2B-C). For each transcript, tumor and normal samples were separately classified at expressing either low or high levels of gene expression based on the mixture component with the largest probability density. This series of filtering steps yielded a set of 3,721 genes that were further filtered, as described below, to identify a set of high-confidence oncogene candidates.

**Figure 1.**
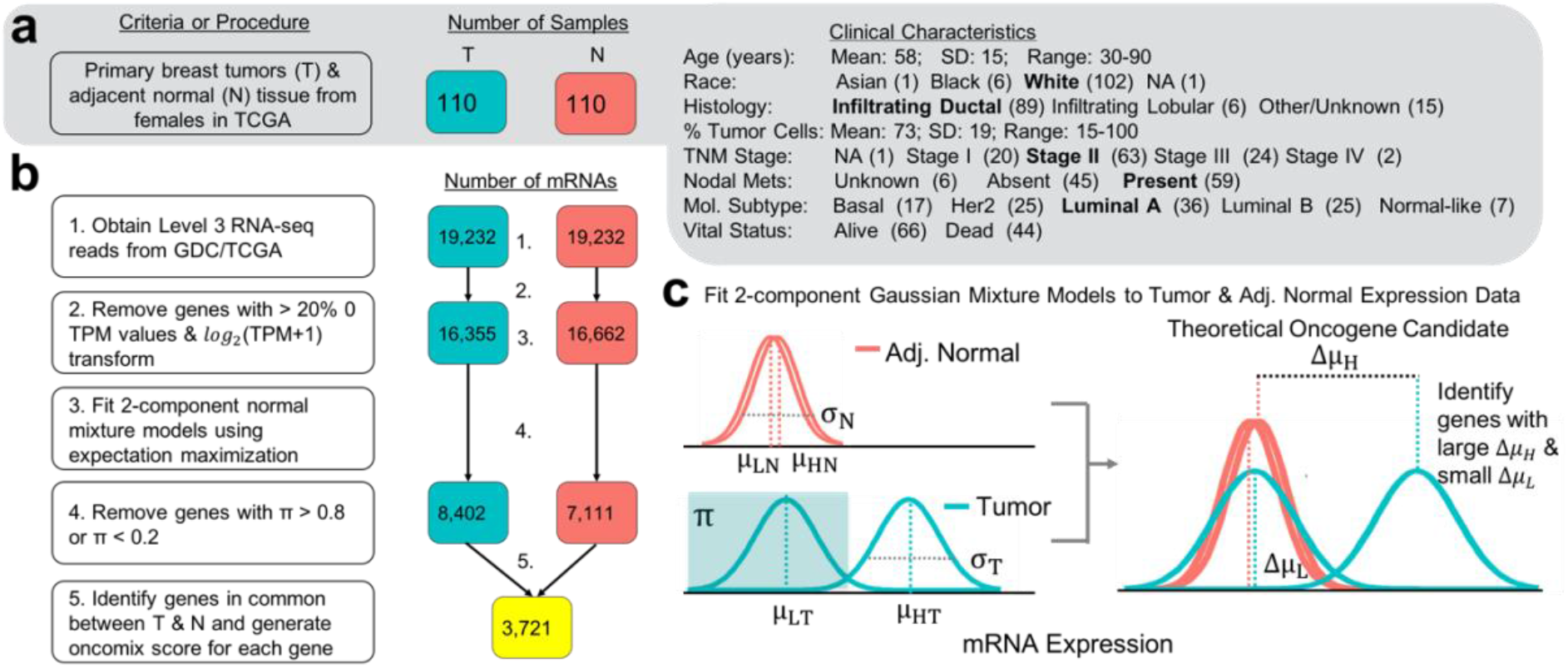
Study design to identify oncogene candidates from breast carcinoma and adjacent normal RNA-sequencing samples. (A) Clinical characteristics of the study cohort of 110 female patients with invasive breast carcinoma. Each of these patients have RNA-sequencing data available from both the primary breast tumor (T) and adjacent normal breast tissue (N). The number of patient samples is indicated within boxes colored either teal for tumor (T) samples, or orange for adjacent normal (N) samples. (B) Workflow of RNA-seq gene filtering based on transcripts per million mapped reads (TPM). The numbered statements on the right reflect the steps used to transform and filter the data for subsequent analysis. The number of genes at each step of the workflow is indicated within the colored boxes (see description in A). (C) An illustration of a two-component Gaussian mixture model (GMM), shown in teal, used to separately fit each gene’s log2(TPM + 1) values for tumor and adjacent normal controls. GMMs yield several distinct parameters; namely, π is the proportion of samples under the Gaussian associated with lower expression values, μL and μH are the means of the curves that fit lower and higher expression values, respectively, and σ is the common standard deviation of the two curves. The additional subscript (T or N) refers to whether the sample parameters are derived from tumor or adjacent normal expression data. Note that the threshold between baseline and overexpressed is defined by the boundary set from the mixture models in the tumor samples and is the point at which the probability of a sample belonging to either the low or high expression group is equal to 0.5.

### *Oncomix* identified five genes with an oncogene-like pattern of expression

Our statistical approach, *oncomix*, detects a distinct bimodal pattern of gene expression across tumors. To identify oncogene candidates (OCs) that matched these specific patterns from the total pool of genes, two metrics were derived from the mixture model parameters. First, a selectivity index (SI) (Figure 2A) distinguishes those genes that are overexpressed in a clearly defined group of patient tumors. A threshold of SI > 0.99 was set based on the observed distribution of the SI values. Examination of the gene expression data from known oncogenes (discussed below, see **Supplementary Figure 1**) with an SI > 0.99 highlighted well-known oncogenes, such as *ERBB2*, in breast cancer. The SI was used in combination with other mixture model parameters to calculate the *oncomix* score, which ranks genes based on their similarity to a theoretically ideal oncogene (Figure 2B). The distribution of expression levels for the five genes with the highest *oncomix* score each demonstrate a clear and distinct subgroup of tumors that overexpress each gene (Figure 2C).

**Figure 2.**
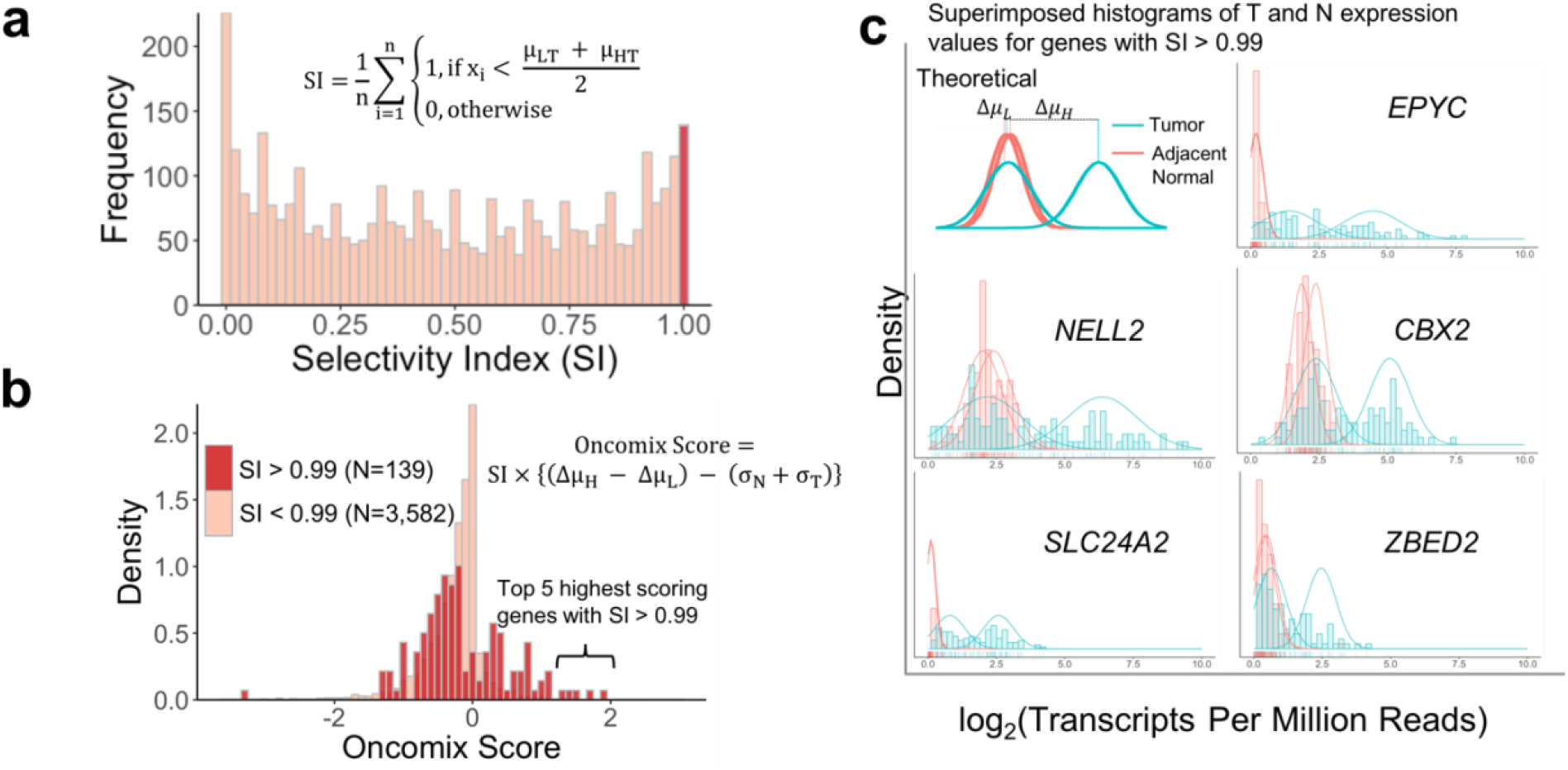
Identification of oncogene candidates using RNA-sequencing data from primary invasive breast carcinomas and adjacent normal breast tissue. A) The distribution of selectivity indices across the 3,721 genes filtered from Figure 1 is shown. The equation for the selectivity index for a gene with adjacent normal and tumor expression values is displayed and is defined in detail in the methods section. B) The distribution of the *oncomix* scores separated by genes with an SI above and below 0.99. Larger *oncomix* scores correspond to genes that more closely resemble the profile of a theoretical oncogene candidate. C) Superimposed histograms of expression values from tumor (teal) and adjacent normal (red) samples for the 5 genes with the highest *oncomix* score and a selectivity index greater than 0.99. The best fitting mixture model is shown for each selected gene. The HUGO gene symbol for each gene is displayed for each histogram. A theoretical model for an ideal oncogene candidate is shown in the upper left and includes some of the summary statistics that were used to compute the *oncomix* score. The y-axis represents density and the x-axis represents log2(TPM + 1) reads. Abbreviations: T = primary breast tumor, N = adjacent normal breast tissue, TPM = Transcripts Per Million reads.

A literature search of the 5 OCs discovered by *oncomix* revealed that oncogene-like features have previously been linked to two of these genes (Table 1, genes in bold). Chromobox 2 (*CBX2*) and neural EGFL like 2 (*NELL2*) have been shown to promote invasion, metastasis, or cell division in a variety of *in vivo* and *in vitro* models of cancer. For example, the gene *CBX2* was recently shown to be highly-expressed in both androgen-independent, late stage prostate cancers (PrCa) and distant PrCa metastases^12^. *CBX2* is a member of the polycomb repressive complex (PRC), and expression of this gene and its protein product is negatively associated with breast cancer survival^13,14^. In addition, *NELL2* is a neural cell growth factor whose expression is positively regulated by estrogen and that promotes invasion of breast cancer cells^15,16^. The sympathetic nervous system has also been shown to promote breast cancer metastasis from primary tumors^17^. These results lend support to the premise for our method, which models population-level patterns of gene expression in subgroups of patients to identify oncogene candidates.

**Table 1.** List of oncogene candidate function and comparison with current differential expression approaches. Each oncogene candidate is represented by a row. Columns indicate the molecular features or function of each gene. Yellow background: A rank-based comparison between the *oncomix* score, limma’s p-value, and mCOPA’s fold change is shown. Genes with a selectivity index > 0.99 were ranked according to the *oncomix* score. A limma rank of 1 is assigned to the gene that was most differentially expressed (ie has the lowest p-value) between tumors and adjacent normal samples, and a limma rank of 7,889 is the lowest possible rank and indicates the gene that was least differentially upregulated in tumors relative to normal tissue. mCOPA identified 2105 genes that contained overexpressed outliers after selecting genes that had at least a log_2_(fold change) > 2 between tumor and normal samples at the 70^th^, 80^th^, or 90^th^ percentile. Genes were ranked according to log_2_(fold change). NA indicates that the gene was not selected by mCOPA.

| Gene symbol | Function (NCBI gene summary) | Chromosome | Oncomix score/ Rank | Limma Rank (out of 7,889 upregulated genes) | mCOPA Rank (out of 2,105 ranked genes) |
| --- | --- | --- | --- | --- | --- |
| EPYC | Member of the small leucine-rich repeat proteoglycan family | 12q21.33 | 1.88 / 1 | 280 | NA |
| **NELL2** | **Neural epidermal growth factor-like like protein 2** | 12q12 | 1.70 / 2 | 2250 | NA |
| **CBX2** | **Member of polycomb repressive complex** | 17q25.3 | 1.47 / 3 | 775 | NA |
| SLC24A2 | Member of calcium/cation antiporter superfamily of transport proteins | 9p22.1-p21.3 | 1.41 / 4 | 148 | NA |
| ZBED2 | zinc finger BED-type containing 2 | 3q13.13 | 1.29 / 5 | 1605 | 379 (best out of 3 thresholds) |

### *Oncomix* recovered a subset of existing oncogenes that are overexpressed in a subset of tumors

While *oncomix* was primarily intended to discover novel oncogenes, it was also imperative to evaluate whether our method could recover any well-established oncogenes. To do this, all Tier 1 oncogenes were used from the Cancer Gene Census (CGC) database (196 genes)^18,19^, a collection of genes with mutations that are causally associated with cancer derived from all tumor types. Of the 196 Tier 1 oncogenes from the CGC, nine genes (4.5%) had an SI > 0.99 and an *oncomix* score > 0 (**Supplementary Figure 1**). The gene expression distributions of these nine genes in the matched tumor-normal samples from the TCGA breast cancer patients showed that most of these distributions contained a subset of tumors that overexpressed the given gene relative to normal tissue (**Supplementary Figure 1**). Of these nine genes, five (*HOXA13, TAL2, SOX2, HOXD13*, and *SALL4*) are transcription factors that help govern embryonic mammalian development and are transcriptionally silent in most adult tissues^20–23^ (**Supplementary Figure 2**). We conclude that our approach successfully identified a small subset of known oncogenes whose function may be mediated through gene overexpression.

### The oncogene candidates identified by *oncomix* represent a unique set of genes that are not reliably detectable by existing approaches

For an oncogene candidate to be detected by *oncomix*, a gene must exhibit a specific expression profile that demonstrates overexpression in a subgroup of cancer patients (Figure 1C). To test whether genes identified by *oncomix* could be identified by existing approaches, we compared our results with those obtained by two other methods to find potential tumor regulators. Limma is a widely-used method to identify differentially-expressed (DE) genes through a regularized Student’s two sample t-test and assumes the presence of a single mode of expression^24^. None of the genes identified by *oncomix* fell within the top 2% of genes ranked by limma (Table 1 and Methods). In addition, benchmarking was performed against mCOPA, a method that ranks a subset of genes based on meeting a fold change threshold between pre-specified percentiles from expression profiles in tumor and normal samples^7^. mCOPA ranked only one out of our five identified OCs, even after pre-specifying three different percentiles (see Methods). We conclude that our method detects unique genes with established roles in oncogenesis and metastasis for a subset of patients, and that these genes are not detectable using existing DE methods that compare tumor and adjacent normal samples.

### Tumors that overexpress *CBX2* manifest transcriptome-wide changes in the expression of cancer-relevant pathways

Oncogenes are often members of molecular signaling pathways and can drive changes in cellular processes, such as cell proliferation, that promote carcinogenesis. Therefore, we sought to determine whether tumors that overexpressed an OC harbored carcinogenesis-related transcriptional changes relative to tumors that did not overexpress a given OC. For each OC, patients were classified into two groups based on whether their tumor overexpressed the OC or not. Differentially-expressed genes between these two groups were identified using limma to determine which genes were up or down-regulated relative to tumors that overexpressed each OC (q < 0.0001 and log_2_(fold change) > 1), see **Supplementary Table 1**). The overexpression of two of the OCs, *EPYC* and *NELL2*, were associated with minor changes in the cancer transcriptome (≤ 5 differentially expressed transcripts). Among the remaining 3 OCs, there were > 95 differentially expressed transcripts. Notably, *CBX2* was the only OC that had more than one differentially-expressed gene that was downregulated, which is consistent with its role as a member of the PRC.

To characterize the genes that were differentially-expressed in tumors that overexpressed each OC, a pathway overrepresentation analysis (POA) was performed using a stringent threshold (see Methods). For all 5 OCs, no gene sets were enriched when examining only the genes downregulated in tumors that overexpress the OC. Significantly enriched gene sets were present for upregulated genes among the OCs *CBX2, SLC24A2*, and *ZBED2*. Genes upregulated in tumors that overexpressed *SLC24A2*, a gene that encodes a solute carrier protein, were overrepresented in the epithelial-mesenchymal transition pathway, a process critical to the metastasis of epithelial cancers^25^. Though *SLC24A2* has not been directly implicated in cancer before, solute carrier proteins have been purported to contribute to cancer through altered energy metabolism^26^. Genes upregulated in tumors that overexpressed *ZBED2* were enriched in immune-related processes, which could be related to immune cell proportion differences between tumors.

Genes upregulated in tumors that overexpressed *CBX2* were enriched in transcripts that map to genes involved in cell cycle-related and proliferation pathways (Figure 3 and Table 2). These results are consistent with previous results that showed differential expression of cell cycle-related pathways following siRNA-mediated *CBX2* silencing in prostate cancer cells^12^. *CBX2* overexpression was associated with the upregulation of genes mapped to genes such as *KIF2C* (log_2_(fold change) = 1.45; q = 1.32×10^−6^), a member of the kinesin family of proteins that are important for mediating microtubule dynamics during mitosis^27^ (**Supplementary Table 2**). The *KIF2C* gene has been demonstrated to be regulated by EZH2, the catalytic subunit of the polycomb repressive complex (PRC) 2, in the context of melanoma, which supports our findings of a link between the *CBX2*, a member of the PRC1 complex, and *KIF2C* expression^28^. These analyses demonstrate that 3 out of the 5 genes identified to be overexpressed in a subset of patient tumors may alter the breast cancer transcriptome in a biologically plausible manner.

**Figure 3.**
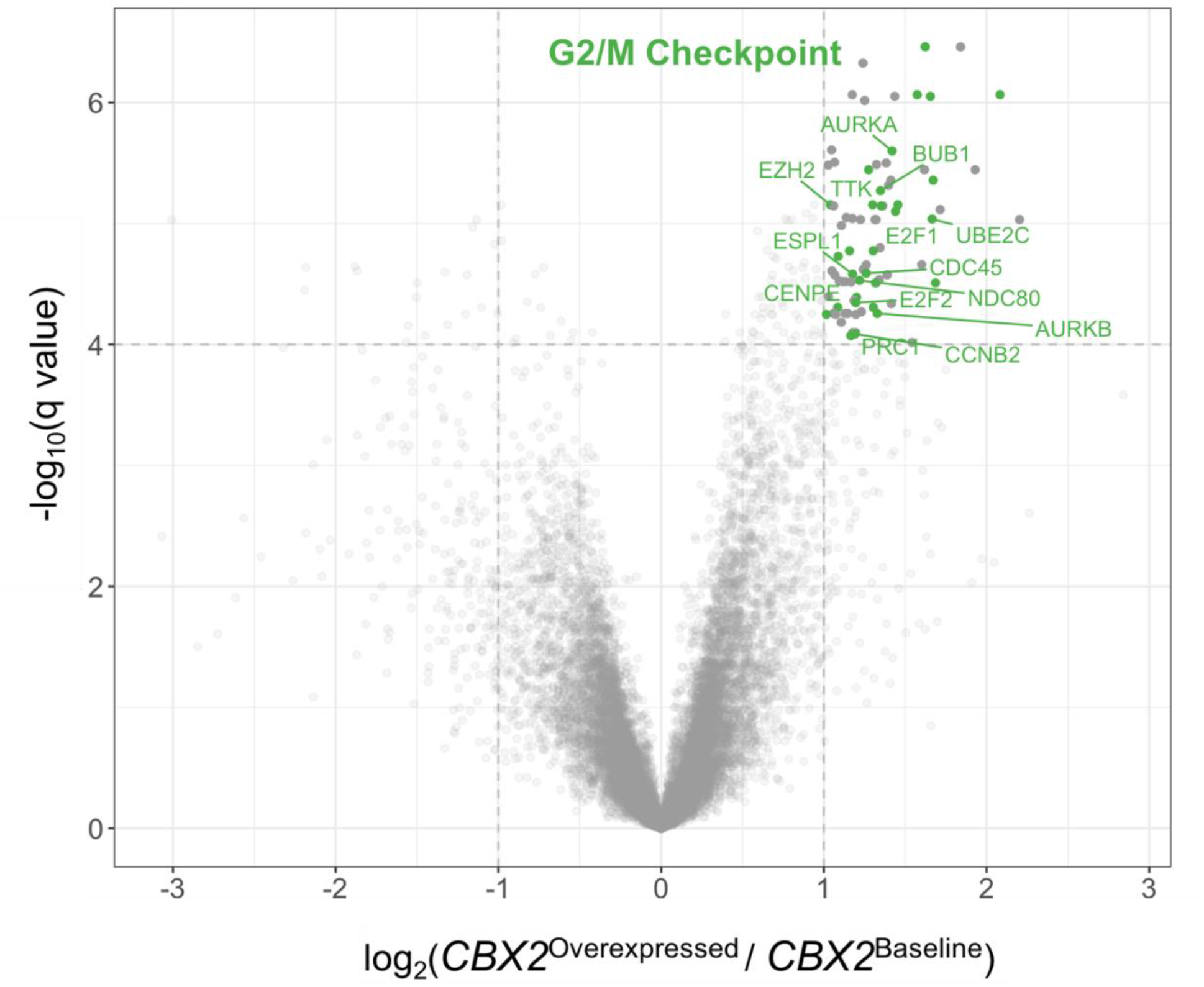
Enrichment of cell cycle processes in upregulated genes within primary breast tumors that overexpress *CBX2* mRNA. A) Volcano plots show 16,157 genes that were tested for differential expression (q < 1×10^−4^ and log2(Mean Fold Change) > 1) between breast tumors that do versus do not overexpress *CBX2*. The mean expression value of a gene in tumors that overexpress CBX2 is denoted as “CBX2^Overexpressed^”, while the mean expression value of a gene in tumors that do not overexpress CBX2 is denoted as “CBX2^Baseline^ “ Significantly upregulated genes (demarcated by the grey dotted lines) within the Hallmark G2/M checkpoint pathway (MSigDB ID: M5901) are highlighted with green points. HUGO Gene Nomenclature Committee (HGNC) symbols are listed for select genes within this pathway. Other genes that are significantly upregulated in *CBX2*^Hi^ tumors are shown in dark grey. q-values were adjusted for multiple testing using the Benjamini-Hochberg method.

**Table 2.**
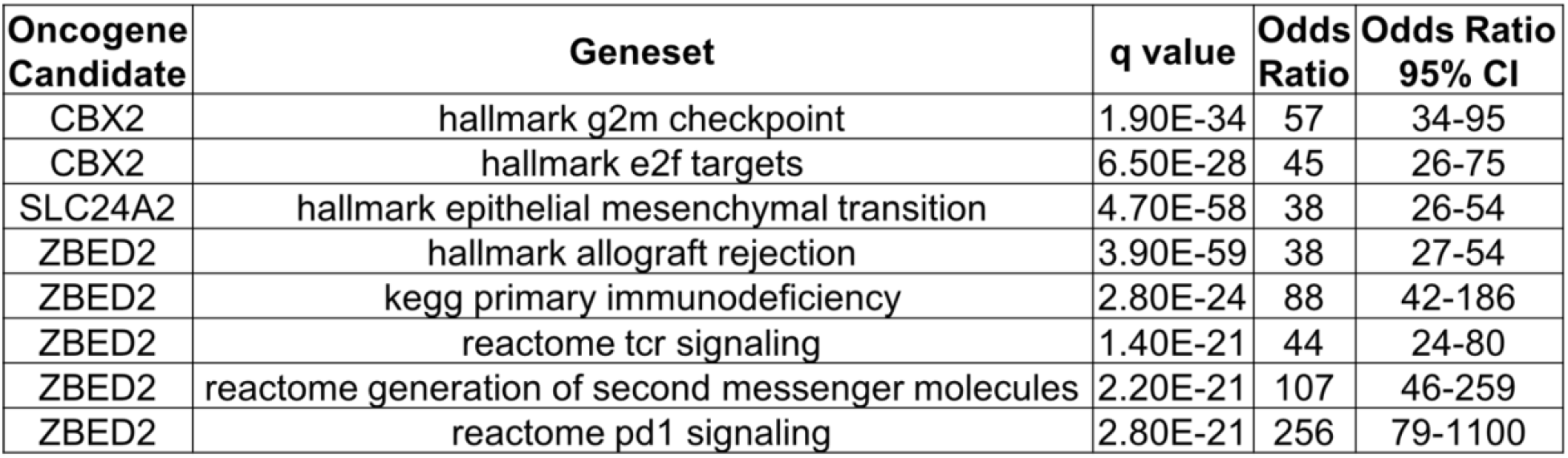
Gene set enrichment from upregulated genes in breast tumors that overexpress a given OC. Three OCs had significant enriched pathways following gene set enrichment performed using Fisher’s exact test. Pathways are shown as rows. Pathways that have an odds ratio with a lower bound 95% CI > 20 and a Benjamini-Hochberg adjusted q-value < 1×10^−20^ are shown and are ranked, from top to bottom, by decreasing odds ratio within each OC. Genes that are differentially expressed in tumors that overexpress *CBX2* (vs tumors that do not overexpress *CBX2*) and found within the Hallmark G2/M checkpoint pathway are highlighted in the volcano plot shown in Figure 3.

| Oncogene Candidate | Geneset | q value | Odds Ratio | Odds Ratio 95% CI |
| --- | --- | --- | --- | --- |
| CBX2 | hallmark g2m checkpoint | 1.90E-34 | 57 | 34-95 |
| CBX2 | hallmark e2f targets | 6.50E-28 | 45 | 26-75 |
| SLC24A2 | hallmark epithelial mesenchymal transition | 4.70E-58 | 38 | 26-54 |
| ZBED2 | hallmark allograft rejection | 3.90E-59 | 38 | 27-54 |
| ZBED2 | kegg primary immunodeficiency | 2.80E-24 | 88 | 42-186 |
| ZBED2 | reactome tcr signaling | 1.40E-21 | 44 | 24-80 |
| ZBED2 | reactome generation of second messenger molecules | 2.20E-21 | 107 | 46-259 |
| ZBED2 | reactome pd1 signaling | 2.80E-21 | 256 | 79-1100 |

### Prediction of OC overexpression reveals that molecular features are more influential than clinicopathologic features

We next sought to identify the biological and clinical features that could contribute to the overexpression of the 5 identified OCs in a subset of breast tumors. The predictor variables used in the regularized multiple logistic regression model represented four broad categories: DNA methylation, expression and copy number, clinicopathologic, and technical variables (see **Supplementary Figure 4** for datasets and processing information and **Supplementary Figure 5** for a model-fitting schematic). For two out of the five OCs, including *CBX2*, intronic methylation was the most predictive covariate. In addition, the molecular subtype, as inferred using Absolute Intrinsic Molecular Subtyping (AIMS) method^29^, was strongly associated with OC overexpression, though not to the same extent as intronic DNA CpG methylation (two-way ANOVA, F(1, 107), AIMS: P-value = 8.9×10^−4^, intronic CpG methyl: P-value = 1.4×10^−9^) (**Supplementary Figure 6**). Relative to the molecular variables, clinicopathologic characteristics, such as cell subtype composition, patient age, and the presence of metastases, were weakly associated with OC overexpression, indicated by the lighter colors and absent within-cell numbers in Figure 4A.

**Figure 4.**
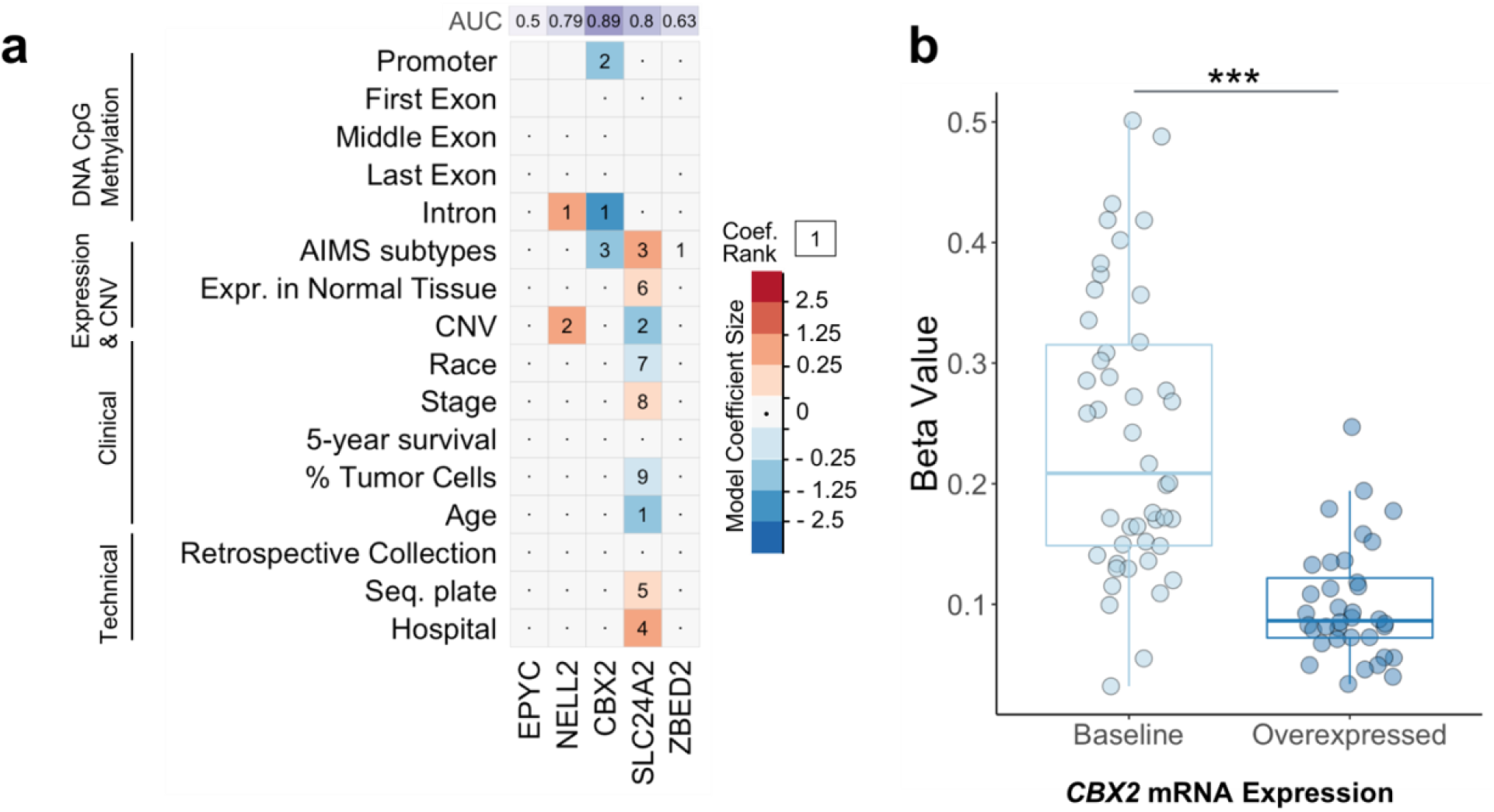
Multi-omic prediction of oncogene candidate mRNA overexpression in breast tumors. A) Visualization of model coefficient selection after regularized logistic regression on binarized (baseline or overexpressed) oncogene candidate mRNA expression levels in breast tumors. Deep blue squares indicate variables that contribute greatly to the prediction of the baseline expression state, while deep red squares indicate variables that contribute greatly to the prediction of the overexpressed state. The numbers in each cell indicate the rank of the absolute value of a coefficient relative to all other coefficients for that model, where 1 is the largest model coefficient. Variables not selected as part of the model are indicated with an interpunct (·). Blank cells indicate missing data for a given model. Each model was used to predict whether a sample overexpressed a given OC or not. These predictions were used to generate receiver operating curves, from which the area under the curve (AUC) was derived (top row, purple background). B) Association of *CBX2* overexpression with DNA methylation beta values for the highest ranking logistic regression coefficient (an intronic CpG locus). DNA methylation values are grouped by level (either baseline or overexpressed) of *CBX2* mRNA expression in tumors. Statistical testing was performed using the Wilcoxon rank-sum test (*** = p < 1×10^−8^).

To validate the utility of the logistic regression models, each model was used to predict the probability of each patient overexpressing the OC in the dataset given her individual features. An area under the curve (AUC) value was generated for each of the five models that predicted overexpression of each OC (Figure 4A, top panel). AUC values greater than 0.8 suggest an excellent fit, while values between 0.7-0.8 suggest a good fit^30^. Models for two out of the five OCs, including the model for *CBX2*, had an AUC greater than 0.8. Furthermore, the distribution of CpG beta values for the single most influential covariate, a CpG located within the 2^nd^ intron of *CBX2*, showed a clear reduction in DNA methylation (P-value < 1×10^−8^, Wilcoxon rank-sum test) in breast tumors that overexpress *CBX2* (Figure 4B). These analyses demonstrate that OC overexpression is strongly associated with molecular covariates, particularly DNA CpG methylation.

### Low levels of DNA CpG methylation in breast cancer cell enhancers is associated with overexpression of *CBX2*

Prior evidence implicates *CBX2* in promoting prostate cancer metastasis^12^ and therefore, clarifying the molecular mechanisms that drive *CBX2* overexpression in the context of this breast cancer cohort has clinical significance for identifying therapeutic targets. Based on the integrative logistic regression model, it was intriguing that against all other potential variables, a single CpG locus within the second intron of *CBX2* was the most predictive factor for overexpression of *CBX2*. This predictor had the largest magnitude across all beta coefficients across all five OCs, indicating that it was more influential than any of the other clinical or genetic factors (Figure 4A-B).

To investigate the relationship between DNA CpG methylation and the functional regulatory elements at the *CBX2* locus in greater depth, a subset of histone and transcription factor ChIP-seq peaks from MCF-7 breast carcinoma cells in ENCODE were overlapped with CpG sites within the *CBX2* locus (see Methods for details). Though ChIP-seq data were not available from the primary tumors themselves, MCF-7 cells represent a valuable model to interpret our results because these cells were derived from a Luminal A breast tumor from an elderly Caucasian woman, a characteristic that demographically matches the profile of many of the patients in the TCGA study. The most significantly differentially methylated CpG locus is found within the second intron of the *CBX2* gene (Figure 5). This CpG site overlaps with two different enhancer marks (H3K4me1 and H3K27ac), promoter marks (H3K4me2 and H3K4me3), transcriptional activation (H3K9ac) and transcriptional elongation (H4K20me1) marks, as well as with a transcription factor, JunD, that promotes cancer cell proliferation^31^. In addition, H4K20me1 is absent in the first exon and promoter region, suggesting that transcription of this gene may begin or be regulated through interactions within the 2^nd^ intron. The overlap between this differentially methylated CpG locus and a JunD binding site raises the possibility that DNA methylation in an active regulatory region regulates JunD binding, possibly through a binary mechanism that regulates gene expression in either a baseline or overexpressed state.

**Figure 5.**
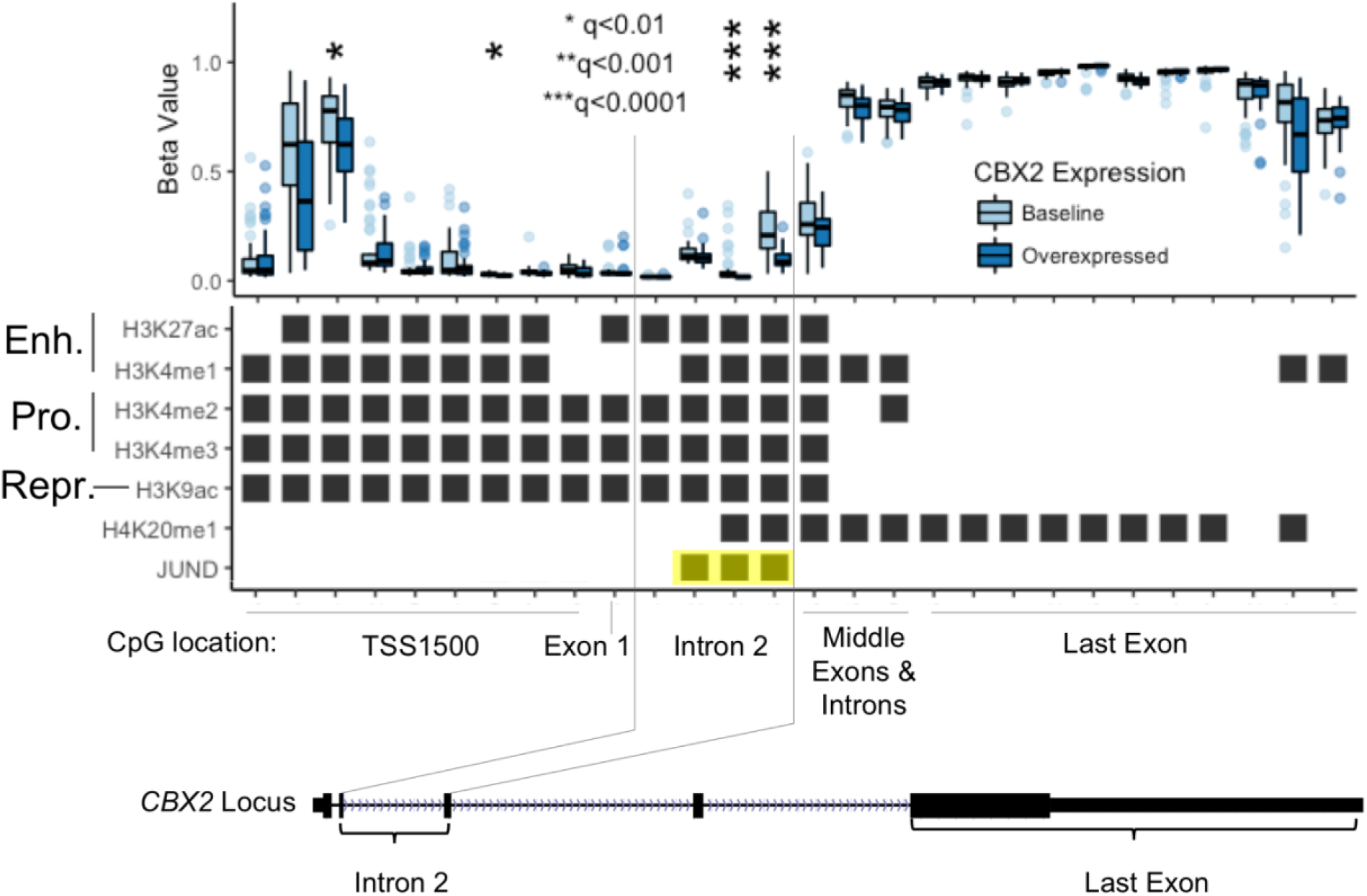
Colocalization of histones and transcription factors with CpG sites that predict overexpression of *CBX2*. (Top) Paired boxplots showing the CpG methylation beta values, which range between 0-1, at each of 28 individual CpG loci for tumors that express baseline levels of or overexpress *CBX2*. (Middle) Each row of the black-and-white matrix represents 1 of 7 different ChIP-seq experiments from MCF7 cells in which a direct overlap (black squares) between a CpG site and a ChIP-seq peak was identified. These 7 ChIP-seq experiments were manually selected for purposes of interpretability from 14 ChIP-seq experiments that overlapped with the *CBX2* locus. The chromatin type or transcription factor is listed along the left-hand side of the matrix, and major chromatin features, such as enhancers (Enh.), promoters (Pro.), and repressive (Repr.) marks, are indicated in large text. Each of the 28 columns represents a different CpG locus within the gene body of the *CBX2* gene (defined as the beginning of the TSS1500 to the end of the 3’ UTR). The model coefficient with the largest absolute value is shown adjacent to the rightmost thin black line. (Bottom) The two thin black lines demarcate the position of the 4 CpG sites within intron 2 and indicate the physical position of these intronic CpG sites within the *CBX2* locus. Additional regions within the *CBX2* gene (length = 11,352 bases, including the TSS1500) are annotated in the gene model, which was obtained from the UCSC genome browser. Asterisks represent q values from a Wilcoxon rank-sum test between the beta values at each of the 28 loci. *** = q < 0.0001, ** = q < 0.001, * = q < 0.01.

### *CBX2* is overexpressed in aggressive breast carcinomas and is associated with poor survival

*Post hoc* statistical testing from the logistic regression model from Figure 4 revealed a significant positive relationship between the aggressively of the Absolute Intrinsic Molecular Subtyping (AIMS) breast tumor subtype and the proportion of patients who expressed *CBX2* within each subtype (multinomial exact test, two-sided P-value = 1.149 × 10^−7^) (**Supplementary Figure 6**)^29^. Expression of *CBX2* is not part of the mRNA expression-based AIMS classification scheme, which highlights the potential utility of *CBX2* in the identification and molecular subtyping of aggressive breast tumors. This is the first report using RNA sequencing data to show that *CBX2* is enriched in basal and Her2^+^ tumors, and our result is supported by a previous study that also found increased *CBX2* expression in basal breast tumors in a microarray mRNA breast cancer dataset^13,32^. We propose that *CBX2* mRNA expression may therefore serve as a marker of aggressive breast cancer subtypes.

### *CBX2* overexpression is associated with poor survival in the TCGA breast cancer cohort

In addition, we searched for differences in survival for patients based on levels of *CBX2* expression (either baseline or overexpressed). A trend toward poorer survival in patients whose tumors overexpressed *CBX2* was detected, though this difference was not statistically significant (q = 0.08, log-rank test). However, survival differences between tumors that overexpressed *CBX2* versus those that did not were also examined in the entire TCGA breast cancer cohort of 1088 patients with available survival data (Figure 6). A significant reduction in 5-year survival in tumors that overexpressed *CBX2* versus those that did not was observed (q = 0.03, log-rank test, Figure 6). This result in consistent with a report that found that high levels of CBX2 protein expression in breast tumors was associated with an increased risk of mortality^14^.

**Figure 6.**
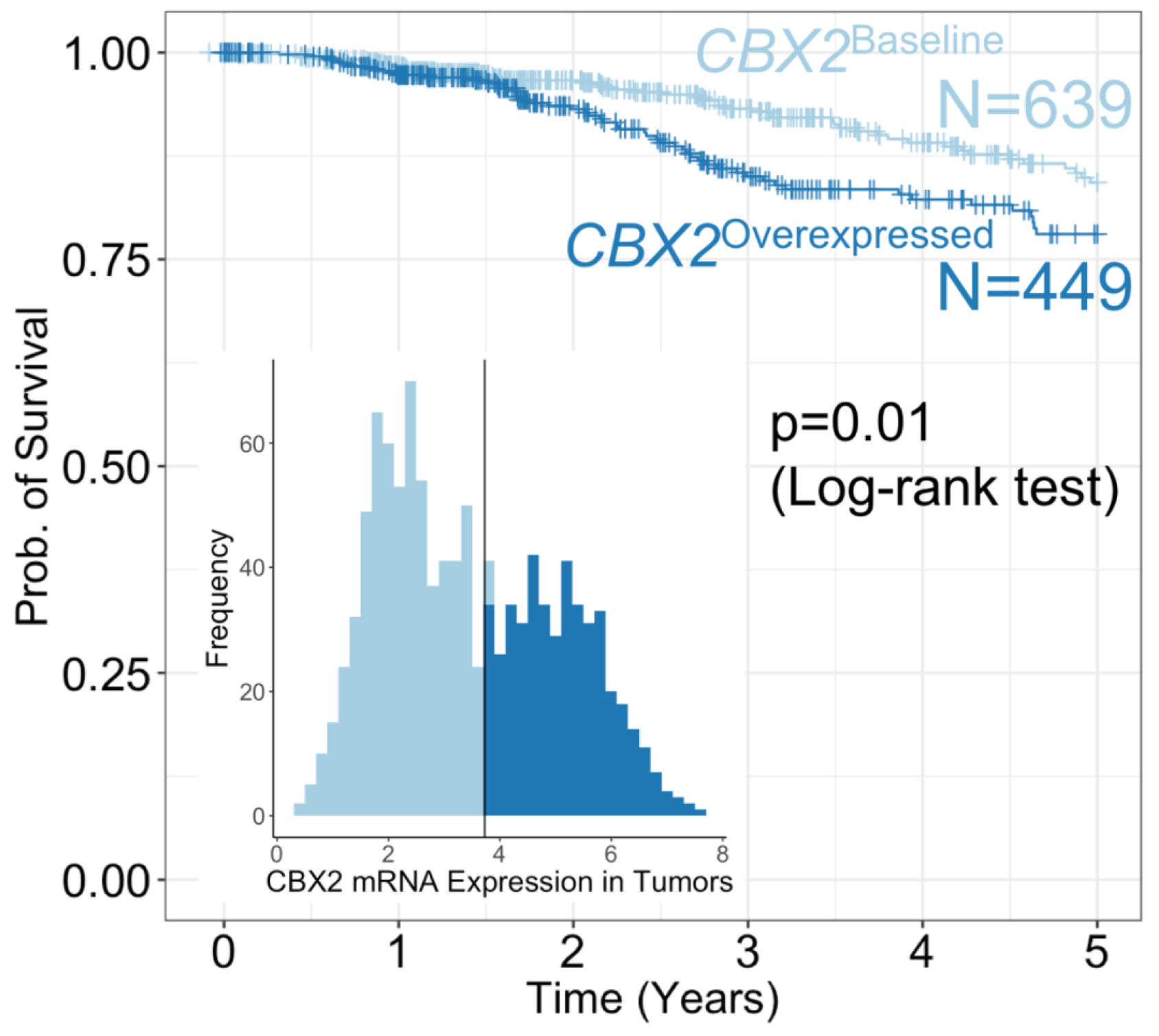
Overexpression of *CBX2* in primary breast tumors is associated with lower rates of survival. A Kaplan-Meier survival curve for 5-year survival rates for 1088 patients with breast tumors from TCGA is shown. Tumors that overexpress *CBX2* are shown in dark blue, and tumors that express baseline levels of *CBX2* are shown in light blue. The tumors were classified using the same boundary that was defined for the original 110 tumor samples. A log-rank test was performed to check for differences in survival between the two tumor types (p = 0.01).

### *CBX2* is expressed at low levels in most adult female tissues

To maximize efficacy and minimize side effects, an ideal drug target needs to be highly expressed in and specific to cancerous tissue, while also expressed at low levels in most other tissues. To examine the expression levels of *CBX2* in normal adult tissues, data from the GTEx portal (https://www.gtexportal.org/home/) was used to examine the expression levels of *CBX2* across 53 normal adult tissues from 8,555 individual samples obtained from 544 human donors. *CBX2* was highly expressed specifically in adult testes and expressed at low levels in virtually all other tissues in both men and women (**Supplementary Figure 8**). Targeted inhibition of *CBX2* may therefore pose a novel therapeutic strategy with minimal side effects on healthy tissue for women whose breast tumors overexpress *CBX2*.

### *CBX2* siRNA knockdown slows the growth of breast cancer cells

Though prior associative computational studies suggest that *CBX2* is linked to breast cancer^13^, no study has experimentally demonstrated a role for *CBX2* in breast cancer. To investigate the role of *CBX2* in breast cancer, we performed genetic knockdown of *CBX2* in MCF7 cells. We observed that adherent MCF7 breast cancer cells grew more slowly following *CBX2* siRNA knockdown relative to a scrambled siRNA control (Figure 7, three-way ANOVA, P-value = 7.0×10^−7^). Furthermore, the number of non-adherent cells was not significantly different between the two siRNA treatments (three-way ANOVA, P-value = 0.08), which suggests that cells either divide more slowly or undergo senescence following *CBX2* siRNA transfection. These results suggest that *CBX2* is involved in regulating the growth of breast cancer cells and that inhibition of *CBX2* function may serve as a therapeutic strategy to slow the rate of breast cancer cell growth.

**Figure 7.**
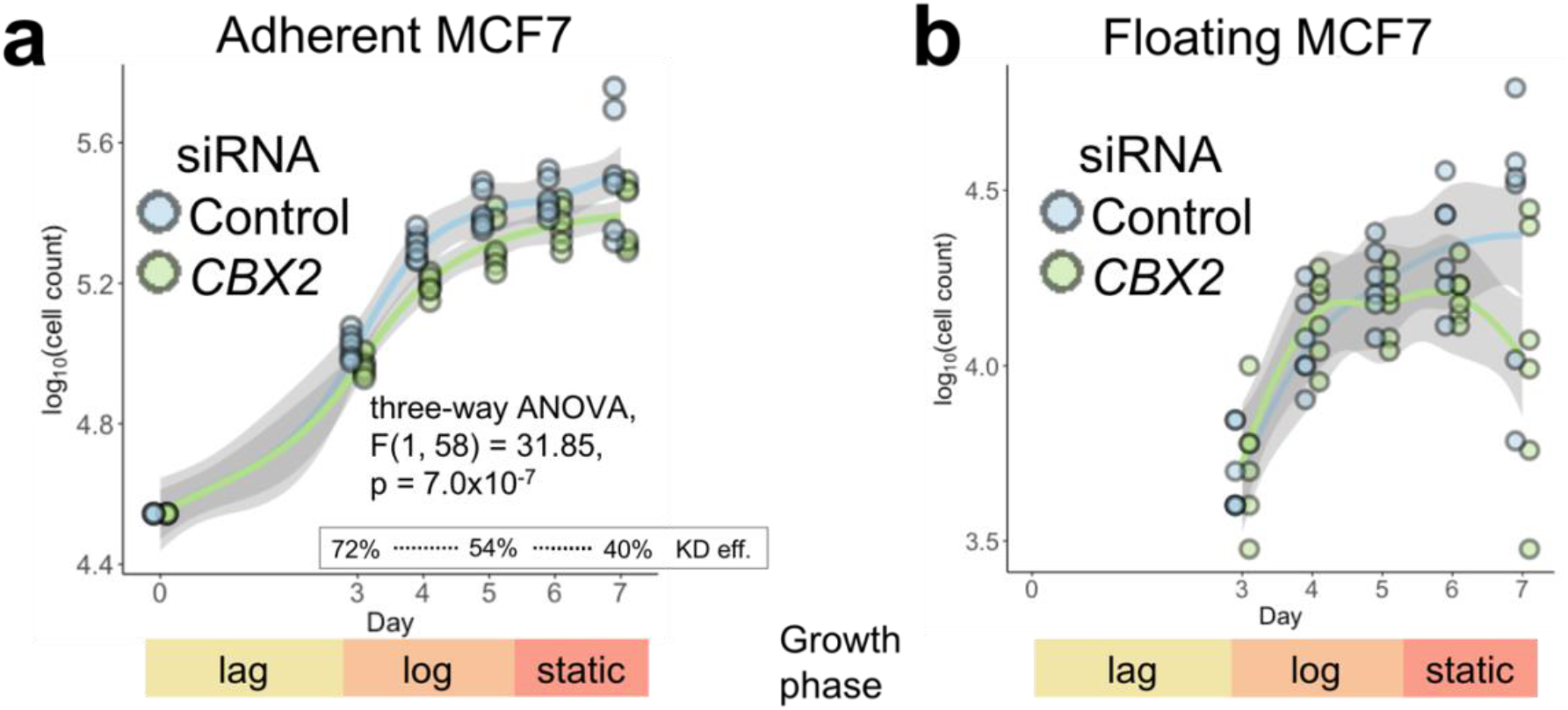
Genetic knockdown of *CBX2* impedes breast cancer cell growth. The cell growth rate for MCF7 breast cancer cells was calculated over a 7-day period following transfection of anti-CBX2 or scrambled siRNA. Both the adherent (alive, panel A) and floating (mostly dead, panel B) fractions of cells were counted. Each point represents one cell count from one of three biological replicates, each with two technical replicates. The 3 growth phases are depicted underneath each plot. KD eff. = *CBX2* knockdown efficiency.

## Discussion

Human breast tumors have a broad array of drivers that modulate growth and metastasis. The identification of additional oncogenic drivers will expand our repertoire of personalized therapeutic targets for breast cancer. Here, we developed a method, termed *oncomix*, that identified oncogene candidate genes (OCs) with known roles in oncogenesis and that unveiled subgroups of patients that overexpress the OC. The value of this tool is made clear by considering *CBX2*, the most promising OC identified, and its implications as a potential drug target for breast carcinoma.

*CBX2* is a gene whose protein product binds to H3K9me3 and H3K27me3 sites with high affinity in mice and forms part of the polycomb repressive complex 1 (PRC1), a multi-protein complex that modifies histones and preserves stemness by silencing lineage-specifying regulator genes in intestinal and embryonic stem cells^33–35^. Our results, which are the first to demonstrate that *CBX2* siRNA knockdown slows breast cancer cell growth, build upon previous studies that showed that *CBX2* siRNA knockdown promotes prostate cancer cell apoptosis^12^. *CBX2* is consistently upregulated in castration-resistant prostate cancer metastases, and its expression correlates with poor patient outcomes in breast and prostate cancer^12–14^. Furthermore, we show that breast tumors that overexpress *CBX2* highly express genes that belong to cell cycle-related pathways. This result is consistent with a prior study which showed that over 500 differentially expressed genes between *CBX2* knockdown and wildtype prostate cancer cells were enriched in proliferation-related processes^12^. Our finding is also consistent with the established role of many oncogenes as drivers of transcriptional alterations within pro-growth signaling pathways^36,37^.

Currently, no successful treatments exist for Her2^+^ and basal breast carcinomas, which are often highly aggressive and disproportionately affect African American and Hispanic women^38^. A therapeutic antibody, trastuzumab, is available as adjunct therapy to treat Her2^+^ breast carcinoma, though a substantial fraction of breast cancer patients develop resistance to trastuzumab^39^. These limitations collectively point to the need to identify new therapeutic strategies to treat these aggressive subtypes of breast carcinoma. Multiple lines of evidence lend support to *CBX2* as a potential drug target against aggressive subtypes of breast carcinoma. First, *CBX2* is expressed at low levels in most healthy adult female tissues, and targeted *CBX2* inhibition may result in fewer side effects than existing treatments. For example, *ERBB2*, the gene that encodes HER2/Neu, is expressed by most tissues in the adult body, which may account for some of the systemic side effects, such as diarrhea, nausea, and cardiotoxicity, seen with the HER2/neu inhibitor trastuzumab. Second, tumors that overexpress *CBX2* also tend to be classified as Her2^+^ or basal, an aggressive subtype against which there are no specific chemotherapeutic interventions, and are associated with poor overall 5-year survival. Third, *CBX2* inhibition via genetic knockdown impedes the growth of breast cancer cells, which suggests that *CBX2* may play an important role regulating breast cancer growth. Fourth, *CBX2* contains a chromodomain that can be pharmacologically targeted, and the crystal structure of *CBX2* was recently solved in complex with a PRC1-specific chromodomain inhibitor, Unc3866^40^. In sum, the results from previous and the current study suggest that *CBX2* is a potential therapeutic drug target in breast cancer.

The identification of a strong association between DNA methylation - a reversible transcriptional regulatory process mediating cellular epigenetic properties - and *CBX2* overexpression suggests that *CBX2* expression may be reversibly regulated to drive important tumor behavior, such as the switch between cell division and metastasis. Prior work suggests a role for *CBX2* overexpression in driving prostate cancer metastasis that was reversible upon siRNA inhibition of CBX2^12^. Metastatic cancer cells undergo reversible changes during the complex processes of extravasation, infiltration, seeding, and proliferation within distant sites, and members of the polycomb complex, such as EZH2, have been associated with metastasis and invasion^41,42^. This apparent plasticity is likely to be governed by epigenetic processes, as opposed to DNA sequence mutations. This is because molecular and cellular plasticity is required to navigate between the dichotomous processes of cell migration, which occurs as tumor cells metastasize to distant tissues, and cell division, which resumes as metastatic tumor cells seed a new site (as reviewed by Tam and Weinberg^43^). The previously published observation that the *CBX2* locus is rarely mutated in human cancers supports the role of *CBX2* in such processes^13^.

Previous studies have found relationships between intragenic enhancers and mRNA expression. For example, the binding of transcription factors to an enhancer within the first intron of *FGFR4*, an oncogene expressed in 50-70% of all pancreatic carcinomas, increases expression of *FGFR4* mRNA^44^. However, intragenic enhancers may also regulate the expression of other genes beyond the gene within which it is located^45^. Furthermore, intragenic enhancers have been found to function as alternative promoters that produce nearly full-length polyadenylated mRNAs with largely unknown functions but that may increase the overall expression of a gene^46^. Furthermore, DNA CpG methylation can directly alter the binding of transcription factors, which supports our hypothesis that CpG methylation may regulate binding of JunD to an intragenic enhancer element in CBX2^47^.

In light of the results identified from *oncomix*, and in combination with existing studies, a conceptual model for the regulation of *CBX2* expression in breast cancer is presented (Figure 8). We propose that *CBX2* is a driver of breast tumor oncogenesis, and that an intragenic enhancer within the *CBX2* locus regulates *CBX2* expression, possibly by acting as an alternative promoter^46^. Within this intragenic enhancer, the binding of a transcription factor that is involved in many cancers, JunD, may be regulated dynamically through DNA methylation. This model is supported by studies that showed JunD binding near proximal promoters and within distal enhancers alters the expression of proto-oncogenes such as Bcl6 and regulators of metastasis such as tissue metalloproteinase^48,49^. In a subset of mostly estrogen receptor-positive breast tumors and within normal breast tissue, the gene expression of *CBX2* remains low, perhaps through active maintenance of DNA methylation. Regulation of *CBX2* expression by DNA CpG methylation may be important for regulating cell division and metastasis, a process that occurs in aggressive breast tumor subtypes (e.g. basal and Her2+) and one that requires dynamic reversibility between cell cycling and cell migration during the epithelial to mesenchymal transition (EMT)^43^. However, the true cause-and-effect relationship between expression and DNA methylation at the *CBX2* locus remains to be fully elucidated.

**Figure 8.**
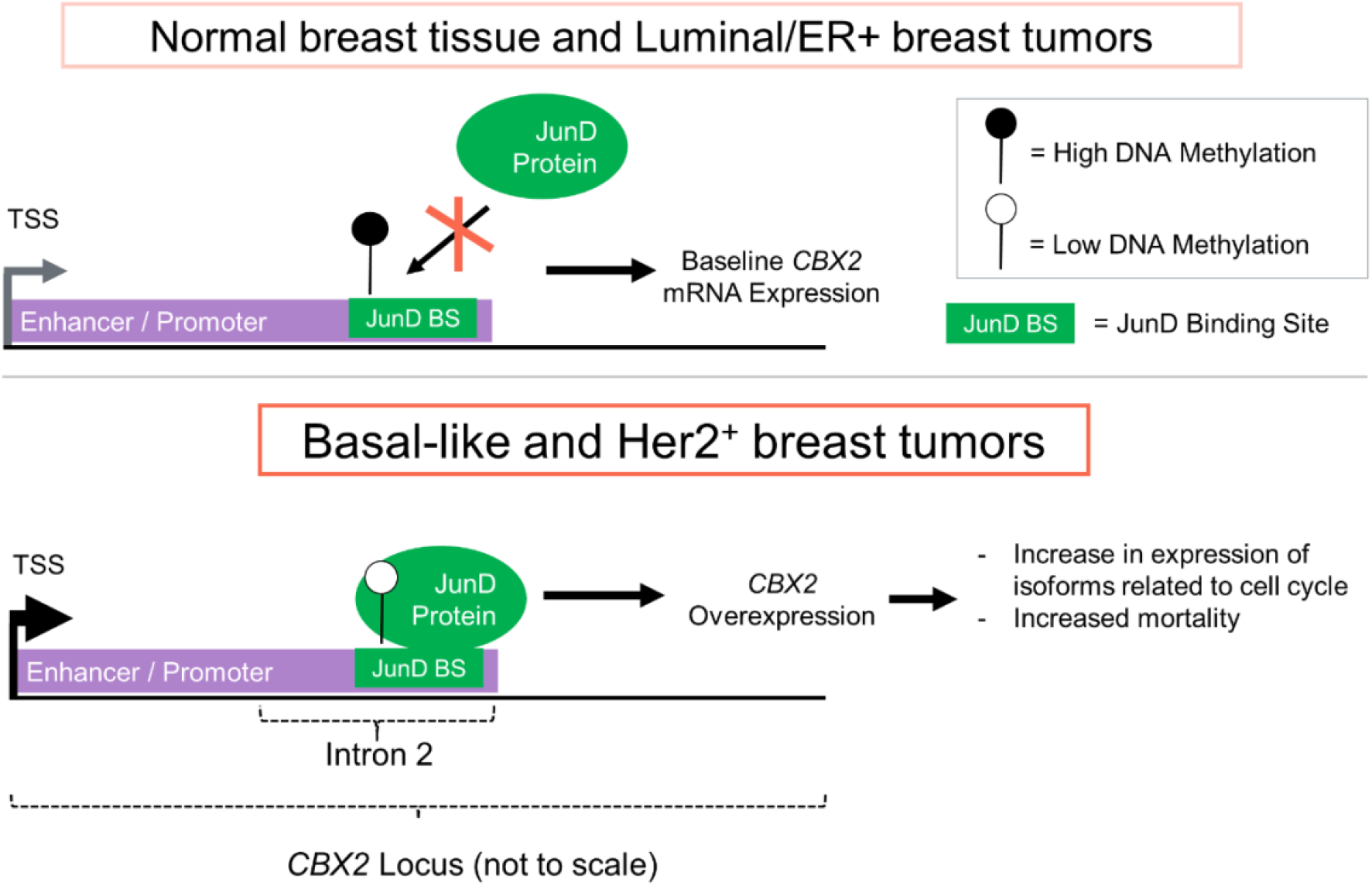
Hypothesized mechanism of the regulation of *CBX2* expression and downstream effects on transcription in breast cancer. The top panel shows a schematic of the molecular basis for *CBX2* expression in normal tissue and in most Luminal/ER+ tumors. Specifically, elevated levels of DNA CpG methylation at an enhancer within intron 2 and at a JunD binding site inhibit the expression of *CBX2*. The bottom panel shows a schematic of *CBX2* overexpression in basal and Her2+ tumors. Low DNA CpG methylation allows for JunD to bind to an intronic enhancer and to increase transcription of *CBX2*, either through interactions with the primary transcriptional start site or through an alternative transcriptional start site.

When comparing the genes identified by *oncomix* versus the other two methods, mCOPA and limma, it was clear that the underlying assumptions made by regarding distributions of the data drive the ranking of the genes. The top five candidates identified by mCOPA and limma highlight how these methods are built to identify genes with specific distributions that deviate from the profile detected by *oncomix* (**Supplementary Figure 3**). Specifically, limma highly ranks genes where the separation between tumor and normal sample means is maximal. mCOPA is designed for the analysis of microarray experiments, is more appropriate for identifying individual outliers, and does not select for genes with visible subsets of patients that overexpress a gene. mCOPA also detects genes that have relatively low variance at the population level for both adjacent normal and tumor tissue. However, *oncomix* is the only method tested that identifies genes for which the tumor samples are grouped into 2 visible clusters (Figure 2C).

Logistic regression modeling proved to be a valuable approach to integrating multiple data types to predict individual OC expression in the breast cancer cohort. However, for two of the OCs, *EPYC* and *ZBED*, it was difficult to train a model that could capture the variance in the observed outcome, suggesting that additional molecular or clinical features not represented in this dataset here may play a role in the regulation of expression of these two genes. To test whether known oncogenic mutations were driving the overexpression of the OCs, a separate analysis was performed to identify the statistical associations between known high-impact oncogenic mutations and the overexpression of each OC (**Supplementary Figure 7**). None of the odds ratios reached statistical significance, though the strongest positive association was between high-impact *TP53* mutations and *CBX2* overexpression (q = 0.053).

In summary, we have identified an oncogene candidate, *CBX2*, based on a theoretical model of identifying subgroups of tumors that overexpress an mRNA gene relative to normal tissue. Computational as well as experimental evidence point to the role of *CBX2* as a regulator of breast cancer cell growth. Our computational method, *oncomix*, is a flexible approach for modeling population-level gene expression data to identify oncogene candidates. Although breast cancer, a well-studied form of cancer, was used as a proof-of-concept example for our method, *oncomix* can be applied to additional types of cancer and to other scenarios where disease-normal pairings are available.

*CBX2* may serve as a potential therapeutic strategy in aggressive breast cancers, due to its low expression in healthy female tissues, available pharmacologic inhibitors, and association with poor survival. Future experimental studies are required to address how DNA methylation within the *CBX2* locus is associated with oncogenic processes such as cell division within both bulk tumor tissue as well as single tumor cells. Our novel approach to identifying OCs through *oncomix* will be particularly useful for identifying regulators of previously unknown tumor subgroups within cancer datasets that include expression levels from hundreds or thousands of patient tumors and their adjacent normal tissue.

## Acknowledgements

Samuel Zimmerman provided advice regarding bioinformatics pipelines and analysis. Raymund Bueno, Shuonan Chen, Ameya Kulkarni, Fabien Delahaye, and Rachel Hazan stimulated helpful discussions and provided critical feedback related to this manuscript. We would like to thank the Molecular Cytogenetics Core at the Albert Einstein College of Medicine - in particular, Dr. Jidong Shan and Dr. Yinghui Song - for assisting with the *CBX2* silencing studies.

## Author contributions

D.G.P. and J.C.M. designed the study and wrote the manuscript. D.G.P. carried out the bioinformatic analysis, created the figures, and interpreted the data. C.M. supervised and performed *CBX2* knockdown experiments. J.M.G and J.C.M. interpreted the data and supervised the project.

## Methods

### RNA Data sources and sample selection

FPKM mRNA-sequencing data from invasive breast carcinoma and adjacent normal controls was downloaded from the Genomic Data Commons web server in January 2018 using the GenomicDataCommons and TCGAbiolinks R packages. RNA from tumors and adjacent normal breast tissue were sequenced by core facilities at the University of North Carolina, Chapel Hill (UNC) on an Illumina HiSeq 2000. Reads were aligned using STAR 2, and BAM files were filtered for quality using samtools and mapped to each gene using HT-seq. Count normalization to FPKM values was performed using custom scripts as described in the GDC workflow (https://docs.gdc.cancer.gov/Data/Bioinformatics_Pipelines/Expression_mRNA_Pipeline/). The FPKM output mapped to 56,963 ensembl gene ids and was converted to transcripts per million (TPM) and subsequently log2(TPM+1) transformed to shrink the numeric range of the data. Genes that contain > 20% zero values were excluded, as genes with many zero values can result in the failure of mixture model algorithms to converge on a set of parameters (unpublished observations). TCGA patient barcodes from the RNA-seq gene level data from both tumors and adjacent normal tissue were intersected, and a total of 110 female patients with RNA sequencing data from both tissue types were selected for further study.

### Supplemental Molecular and Clinical Datasets

All supplemental data discussed in this paragraph was downloaded from GDC servers in January 2018 using the GenomicDataCommons and TCGAbiolinks R packages. 75% (82/110) of tumor samples in this study also had DNA methylation data processed on Illumina 450k arrays that was obtained from the same tumor. The FDb.InfiniumMethylation.hg19 R package was used to obtain 450k CpG coordinates for hg19, which were mapped to hg38 using the rtracklayer R package^50,51^. DNA CpG methylation loci beta values were obtained from Illumina 450k arrays (see **Supplementary Figure 4**). For the logistic regression analysis, only those CpG methylation loci from the TSS1500 to the 3’ UTR within each respective oncogene candidate were used. The TxDb.Hsapiens.UCSC.hg38.knownGene R package was used to obtain the genomic coordinates for each oncogene candidate^52^. Log2 mean segment copy number values for CNV obtained from an Affymetrix 6.0 SNP array were utilized. Clinical data was numerically codified or scaled to within a range of 0-1, and the molecular subtype was inferred from the log2(TPM+1) mRNA expression data from each tumor using the AIMS algorithm^29^.

All 66 transcription factor and histone ChIP-seq data from MCF7 cells with 2 biological or technical replicates was downloaded from ENCODE servers using the ‘rutils’ tool in April 2017. All downloaded data was aligned to hg38, and peaks were called using standard ENCODE processing pipelines^53,54^. For transcription factors, final peak calls were determined and the optimal set of peaks was derived from IDR analysis of biological replicates and pseudoreplicates. For histones, peaks were selected using the narrowPeak algorithm from peak calls that were observed either in both replicates, or in two pseudoreplicates of the pool. All final histone peaks passed an optimal IDR threshold set at 2%. Of the 66 ENCODE data sets, 14 (three transcription factors and 11 histones) overlapped with at least one CpG site within the *CBX2* locus. From these 14 ChIP-seq data sets, seven ChIP-seq experiments were manually selected based on their established association with transcriptional regulation^54^. Final peak lists for each ChIP-seq experiment were overlapped with CpG sites using the GenomicRanges package in R^55^.

### Estimation of mixture model parameters for RNA Seq Data

To investigate whether certain genes expressed in tumors exhibited distinct, clearly separable clusters of gene expression values, a 2-component Gaussian mixture model was fit to each gene across the 110 data points. These mixture models were applied separately for gene expression values from both tumors and adjacent normal samples. For each gene within each group (either tumor or adjacent normal), 4 parameters - namely, the mean of the Gaussian with the lower (μL) and higher (μH) mean, the proportion of samples under the Gaussian with the smaller of the two means (π), and a common standard deviation (σ) - were estimated using maximum likelihood through the well-established method of expectation maximization^56^ (Figure 1C). The variance of the mixture model was set to be equal between the two Gaussians to stabilize the expectation maximization procedure. Each parameter includes an additional letter subscript (“T” or “N”) to denote whether the parameter refers to the model describing the tumor (T) or adjacent normal (N) expression data.

### Selection and filtration of genes

To remove genes with extreme outliers and to allow for sufficient statistical power for downstream analysis, genes with a proportion of low-expression modal membership between 0.2 > π_T_ & π_N_ > 0.8 were selected. Additional filtering of genes was performed as described in Figure 1B. To identify and rank genes whose expression values defined a distinct subgroup of tumors that overexpressed the gene relative to normal tissue, two statistics was derived from the mixture model parameters. The first, termed the selectivity index (*SI*), was used to screen candidate genes with an overexpressed subgroup of tumors and was defined as follows:

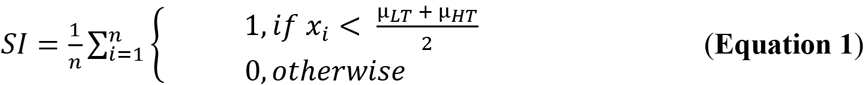

where *n* is the number of paired samples with gene expression values (here, *n* = 110), *x*_i_ is the log2(TPM+1) expression value of the *i^th^* adjacent normal sample, and 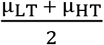 is the boundary, or point of equal probability, between the low and high expression modes of the Gaussians that describe the tumor data. The SI is applied separately to each gene and ranges between 0 and 1, with values closer to 1 indicative of genes that have a subpopulation of samples that are clearly distinct and separable based on the expression values from tumors for a given gene. The SI is unique in that it selects genes that define distinct clusters of tumor samples based on expression values that are separate from and greater than their adjacent normal counterparts as well as from other tumor samples. After visually inspecting the distribution of SI values for all genes (Figure 1A), a conservative SI cutoff of 0.99 was selected.

The second statistic that was developed was termed the *oncomix* score. The *oncomix* score is calculated as a function of the SI (see Equation 1) and the Δ*μ_H_*,Δ*μ_L_*,*σ_N_*,*σ_T_* parameters, as shown below:

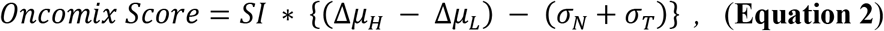

where ΔμH = μHT - μHN and is the difference between the means of the high expression groups of the mRNA values from tumor (μHT) and adjacent normal tissue (μHN). This term, when large, indicates greater separation between the high expression modes of the tumor and adjacent normal populations and would contribute to a larger and more favorable *oncomix* score. The difference between the low expression groups of the tumor (μur) and adjacent normal samples (μLN) was calculated as Δμ_L_ (μur - μLN). This term, when small, indicates a minimal difference between the low expression modes of the tumor and adjacent normal populations and results in a larger *oncomix* score. The *oncomix* score is penalized by the variance of each mixture model (*σ_N_* & *σ_T_*), with larger variances resulting in lower scores. This is because mixture models with large variances reflect an underlying spread in the distribution and provide evidence against the existence of two distinct clusters of tumor expression data, and of a single cluster of normal tissue data.

### Benchmarking oncomix against limma and mCOPA

Differential expression between tumor and adjacent normal samples was performed using limma, an established method for performing a 2-sample t-test in conjunction with empirical Bayes estimation^24^. 16,158 genes that had >20% non-zero values for both tumor and adjacent normal samples were used and ranked using the t-statistic and resulting p value. A ranking of 1 indicates the gene with the smallest p value. Permutation q-values were calculated by uniformly sampling without replacement 1×10^5^ times from a distribution of possible rankings and comparing how frequently the sampled ranking was smaller than the observed rank (see supplemental Rmarkdown file). Expression data for 16,158 genes from 220 paired tumor-adjacent normal samples was used as input into mCOPA. mCOPA requires the manual specification of percentiles and was run three times using the 70^th^, 80^th^, and 90^th^ percentile. The 80^th^ percentile results were displayed in **Supplementary Figure 3**, with the rationale that these would be most consistent with our requirement that at least 20% of samples appear in either the high or the low expression group.

### Differential expression and pathway overrepresentation analysis

Differential expression analyses was performed using limma^24^. The threshold used for differential expression was a Benjamini-Hochberg adjusted q-value of 0.0001 and a log_2_(fold change) > 1 or < −1. Pathway overrepresentation analysis (POA) was performed using 910 gene sets from three well-defined, manually-curated pathway databases - Hallmark^57^, KEGG^58^, and Reactome^59^. POA was performed separately for significantly upregulated and downregulated genes to facilitate interpretability, and a stringent cutoff (q < 1×10^−20^ & OR_95% ci_ > 20) was used to select highly enriched gene sets.

### Multiple logistic regression, variable selection, and coefficient shrinkage using the elastic net

Multiple logistic regression was performed for each OC with binary response variables (normal or overexpressed OC mRNA levels in breast tumors) and complementary clinical, molecular, and pathological datasets were used as covariates (see **Supplementary Figure 4** for datasets and processing information). The output from the logistic regression model provides a weight, in the form of a beta coefficient, that estimates the influence for each predictor on the response variable, which in this case, is the overexpression of the OC. How strong of an influence the predictor has on the response is estimated by the model, as well as the direction of this influence. To prevent model overfitting, the size of the model coefficients, whose effect was assumed to be additive, were regularized using the elastic net penalty and leave-one-out cross validation^60^ (see **Supplementary Figure 5**). The elastic net is a regularization term that shrinks and selects model coefficients to prevent overfitting of data, particularly in settings when there are many predictor variables, and helps account for potential collinearities between covariates^60^. Here, the elastic net was used to shrink and select the model coefficients weights for our logistic regression model, where the binary outcome variable is the level of expression (either baseline or overexpressed) for a given gene. The implementation of the elastic net in the R package ‘glmnet’ was used with an α value fixed at 0.5. The multiple logistic regression model was fit using penalized maximum likelihood through solving the following objective function (Equation 3) using coordinate descent (as implemented in glmnet^61^):

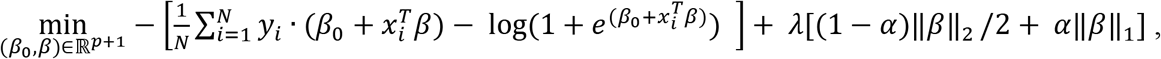

where *β_0_* is the model intercept, *β* is a column vector of regression coefficients, 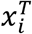 is a row vector of scaled variables (observations) for the *i^th^* individual, *y_i_* is the expression status (either baseline or overexpressed) for an oncogene candidate, *N* is the number of individuals in the dataset (here, N=110). The right half of the objective function (outside of the large brackets) represents the elastic net regularization term. The purpose of this term is to prevent overfitting by selecting and shrinking the *β* coefficients and is particularly useful as the number of variables approaches the number of observations. The objective function is penalized by the size of the beta coefficients. Specifically, ||*ß*||_1_ and ||*ß*||_2_ are L1 and L2 penalty terms on the magnitude and square of the magnitude of the beta coefficients. *λ* regulates the overall size of the penalty term and was selected using leave-one-out cross validation across a grid of *λ* values. The selected *λ* value is associated with the sparsest model that yields a misclassification error (MCE) within 1 standard error of the MCE. If *λ* = 0, then the solution to this problem is equivalent to the estimates obtained by ordinary least squares^60^. *α* is a manually-set tuning parameter that ranges between 0 and 1. When *α* = 1, the regularization term is known as the LASSO (L1 penalty), when *α* = 0, the regularization term is known as ridge regression (L2 penalty), and when 0 < *α* < 1, this regularization term is known as the elastic net. Here, *α* was set to 0.5 for all models. All continuous variables were scaled between 0 and 1, and all categorical variables were coded as binary indicator variables with a separate column per category. A table of all variables used and the method for variable scaling are available in the supplementary RMarkdown file.

### Gene set enrichment analysis

The Hallmark^57^, Kegg^58^, and Reactome^59^ geneset databases were downloaded from MSigDB as GMT files in March 2017^62^. To test whether the differentially expressed genes between tumors that do vs. do not overexpress a given oncogene candidate were overrepresented in any of the 910 genesets obtained from these three databases, a Fisher’s exact test was performed. Genesets that had an odds ratio with a lower bound 95% confidence interval > 20 and a q < 1×10^−20^ corrected using the Benjamini-Hochberg method were selected.

### Code availability

All analysis was performed in the statistical programming language R (version 3.4.3). An HTML document created using knitR and RMarkdown contains the code and workflow for all analysis performed in this study (**Supplementary File 1**). An R package “oncomix” for identifying oncogene candidates in large cohorts of RNA-sequencing data from tumor and adjacent normal samples is available through Bioconductor^63^.

### Data availability

All of the data used in this study, with the exception of the siRNA knockdown experiments, was publicly available and was downloaded from the genomic data commons, Encode, and Gtex databases. Data related to siRNA knockdown experiments are available upon request.

### Statistical analysis

All statistical tests were two-sided unless otherwise noted. All statistical tests were performed in R (version 3.4.3), and implementations of specific statistical tests can be found in **Supplementary File 1**.

### CBX2 siRNA knockdown experiments and analysis of cellular growth rate

MCF7 cells were obtained from ATCC (#HTB-22). Cells were grown in DMEM supplemented with 5% fetal bovine serum and 0.01 mg/ml human recombinant insulin (Sigma) and incubated in 5% CO2/37°C. For silencing of CBX2 the siRNA SMARTpool (L-008357-Dharmacon, Lafayette USA) was used. On-target CBX2 oligonucleotides were used for gene-specific downregulation and same MCF7 cells transfected with the Non-Targeting (Scramble) siRNA Control Pools were used as a reference control for all experiments. SiRNA pools were resuspended using according to the manufacturer’s protocol in RNase-free 1x siRNA Buffer at a final concentration of 20 mM. Cells were transfected using DharmaFECT-4 Transfection Reagent according to the manufacturer’s instructions. After transfection, cells grew for 48 hours before the analysis of specific endpoints.

For the growth curve analysis, MCF7 cells silenced with the siCBX2 SMARTpool and scramble controls were plated at ∼17,000 cells/cm^2^ in 24 well plates, incubated at 37°C for 48 hours and the cell number counted in duplicate every 24 hours for five days. All experiments were repeated three times in independent biological triplicates. MCF7 were routinely analyzed to ensure lack of mycoplasma contamination by DAPI staining. A three-way between-subjects ANOVA without interaction terms was conducted to test the null hypothesis that siRNA has no effect on cellular growth rate. The independent variables, all categorical, were the siRNA, the biological replicate, and the day post-transfection. The MCF7 cell line was authenticated using the GenePrint 24 system (Catalog number B1870, Promega) and analyzed using the GeneMarker 1.75 software (SoftGenetics). Cell line genotypes showed 100% identity to MCF7 cell lines (results available upon request).

### RNA isolation and cDNA synthesis to evaluate CBX2 levels

MCF7 siCBX2 and siScramble were established as described above and plated in 6 well plates at ∼17,000 cells/cm^2^ for 48 hrs. Cells were then analyzed at 72-120-168 hrs post transfection. The cells were then lysed directly on the plate with Qiazol lysis reagent (Qiagen, Valencia, CA) and placed at −80°C until all samples were ready for RNA extraction. Total RNA was isolated using the miRNeasy kit (Qiagen, Valencia, CA). cDNA was reverse-transcribed from 5 μg of total RNA using random primers and SuperScript II Reverse Transcriptase (Invitrogen). *CBX2* and *GAPDH* primers were designed with Primer3 software (sequences listed below). Real-time qRT-PCR was performed using Applied Biosystems Fast SYBR Green Master Mix and the StepOnePlus Real-Time PCR System (Life Technologies Corp., Carlsbad, CA, USA). Data normalization and analysis were performed as previously described (Acosta *et al.*)^64^.

CBX2fw: 5’ - GGCTGGTCCTCCAAACATAA-3’

CBX2rev: 5’ - GCACCTCCTTCTC ATGTTCC-3’

GAPDHfw: 5’ - CCACATCGCTCAGACACCAT −3’

GAPDHrev:5’ - CCAGGCGCCCAATACG −3’

